# Enhanced carbon storage in dissolved organic matter in a future oligotrophic ocean

**DOI:** 10.64898/2026.04.02.716036

**Authors:** Takasumi Kurahashi-Nakamura, Thorsten Dittmar, Adam C. Martiny, Sinikka T. Lennartz

## Abstract

Marine dissolved organic carbon (DOC) is one of the ocean’s largest carbon reservoirs, which persists for millennia and exceeds the carbon stored in terrestrial and marine biomass combined (Hansell et al, 2012; Dittmar et al, 2021; Moran et al, 2022). Yet its response to climate change remains uncertain, largely because the ecological mechanisms behind its microbial degradation are poorly understood (Wagner et al, 2020; Legendre et al, 2015; Lønborg et al, 2020). Here we show how observed global-scale distribution of DOC emerges from bottom-up ecological controls acting on microbial DOC consumers. We combined large-scale metagenomic data (Larkin et al, 2021) with incubation experiments (Hale et al, 2017) to map global patterns of nutrient limitation in heterotrophic microbial communities responsible for DOC uptake. We then integrated a new microbial–DOC component into an Earth system model, which successfully reproduced observed global distributions of nutrient stress and DOC concentrations. Linking carbon and nutrient cycles reveals quantitatively significant consequences for the future ocean. Under a high-emission scenario (SSP5-8.5), the DOC pool is projected to increase by 18–44 gigatons by 2200. This increase, driven by intensified nutrient limitation in surface waters, contributes substantially to the biological carbon pump, accounting for about **∼**30% of the increase in deep-ocean carbon storage associated with particulate organic carbon export. Overall, our results indicate that the marine DOC reservoir is more dynamic than previously thought. Reduced DOC remineralisation in an increasingly oligotrophic ocean constitutes a quantitatively significant negative feedback on centennial timescales.

## 1 Main

The ocean plays a central role in Earth’s climate by absorbing ∼30% of anthropogenic CO_2_ emissions (e.g. Friedlingstein et al, 2025). DOC is a diverse mixture of compounds with turnover times from seconds to millennia and constitutes the ocean’s largest organic carbon pool with ∼700 Pg of carbon (Druffel et al, 1992; Dittmar et al, 2021; Moran et al, 2022). This considerable carbon storage is largely attributed to microbial alteration and diversification of photosynthetic organic compounds that increases their recalcitrance (Dittmar et al, 2021; Jiao et al, 2010), as conceptualised in the microbial carbon pump (MCP) framework (e.g. Jiao et al, 2010).

Understanding the fate of this carbon reservoir under future environmental change is crucial for future projections and carbon management strategies, yet its response remains highly uncertain. A key limitation is the lack of quantitative understanding of the processes that shape basin-scale DOC distributions (Legendre et al, 2015; Wagner et al, 2020; Lønborg et al, 2020). This lack of understanding translates into an insufficient representation of interactions between dissolved organic matter (DOM) and microbes in state-of-the-art Earth System Models (ESMs), our most powerful tools for projecting future climate scenarios. Traditionally, global ocean models–including modern ESMs–represent DOC by partitioning it into reactivity classes that decay via first-order degradation rates (e.g. Bacastow and Maier-Reimer, 1991; Kirchman et al, 1993; Yamanaka and Tajika, 1997; Levine and DeVries, 2024). While these net-removal-rate models successfully reproduce present-day DOC distributions, they lack dynamic responses to environmental shifts that affect microbial activity and DOC turnover (e.g. Levine and DeVries, 2024). Consequently, future DOC projections currently reflect only changes in production, neglecting environmental controls on degradation (Gilchrist, 2022).

The amount of DOC stored in the ocean is determined by the balance of DOC production by phytoplankton, and DOC consumption by heterotrophic microorganisms (Carlson et al, 2024). Nutrient dynamics are central to these microbially mediated carbon fluxes (Romera-Castillo et al, 2016). Primary production, the ultimate DOC source, depends on nutrient availability and is itself affected by the remineralization of organic nutrients (Najjar et al, 1992). Similarly, incubation experiments indicate that heterotrophic DOC degradation is nutrient-limited across major ocean basins, (e.g. Pesant et al, 2015; Clayton et al, 2022; Romera-Castillo et al, 2016; Hale et al, 2017). While metagenomic data from large-scale surveys provide unprecedented insights into the functioning of the oceanic microbiome, they have remained largely untapped for model development purposes. A major obstacle in this respect is linking the genetic potential with macroscopic traits amenable to biogeochemical models. First efforts to integrate such microbial traits–including nutrient requirements (Lennartz et al, 2024) or taxonomy (Zakem et al, 2025)–have improved the reproduction of observed DOC patterns and have demonstrated the potential of using molecular and ecological information for Earth system model development (e.g. Tagliabue, 2023; Saito et al, 2024). Through these modeling efforts, millennial-scale net removal rates of DOC have been linked to rapid microbial processes controlled by macronutrient availability (Lennartz et al, 2024), implying greater temporal variability of the marine DOC inventory than previously thought. First future simulations, in which only the labile fraction dynamically responds to environmental controls, indicates a 6% change in DOC export in a future climate scenario (Flanjak et al, 2025). Yet, no future scenario simulation exists in which the whole DOC pool dynamically responds to changing environmental conditions, which is a prerequisite to assess the future of the global DOC inventory.

Here, we address this fundamental lack of knowledge by combining global metagenomic data and results from incubation studies in an ESM - with and without an explicit formulation of the remineralization of DOC by microbes. In the model version without an explicit microbial representation, DOC fractions decay according to first-order kinetics, as in most established ESMs (e.g. Letscher et al, 2015). First, we show that the geographic pattern of the genetic potential for N- and P-transporters in marine bacteria mirror the nutrient limitation of their growth rates in incubation studies. Building on this, we developed a bidirectionally coupled module for an ESM to assess the basin-scale distribution of DOC, building on Lennartz et al (2024). The dynamical DOM module explicitly depicts the production of DOM through primary production, its degradation by heterotrophic microbes (interactively coupled UVic ESCM–MICDOC model; see Methods), and the dynamics of DOC and dissolved organic nutrients. For the first time, this new framework allows us to explore how microbial nutrient limitation influences DOC dynamics under future climate scenarios, where surface nutrient concentrations are expected to decline significantly (e.g. Chavez et al, 2011; Falkowski, 2012). Our approach reveals that DOC accumulation and turnover are more sensitive to changes in nutrient concentrations than traditional models suggest. Quantifying these dynamics is essential to project the ocean’s carbon storage capacity and its feedbacks in the climate system in future scenarios.

### Limiting factors of microbial carbon consumption

DOC accumulates at the surface to concentrations typically between 45 to 80 mmol m*^−^*^3^, with highest concentrations found in oligotrophic subtropical gyres (Hansell et al, 2021). Its concentration decreases during the overturning circulation, so that the oldest water masses in the Pacific ocean show concentrations around 35 mmol m*^−^*^3^ (Hansell et al, 2012; Hansell, 2013b). This observable DOC represents the fraction not immediately consumed and respired by heterotrophic microorganisms(Hansell, 2013a). Understanding the factors that limit microbial DOC uptake is therefore essential for assessing basin-scale controls on its distribution.

Several lines of evidence demonstrate that the degradation of DOM is regionally limited by the availability of macronutrients such as nitrogen and phosphorus. Numerous nutrient enrichment experiments, in which the heterotrophic community has been incubated in the dark with the addition of nitrogen, phosphorus and/or glucose, have shown that these inorganic macronutrients limit bacterial activity alone or in combination with organic carbon (colimitation) (Carlson and Ducklow, 1996; Caron et al, 2000; Chin-Leo and Benner, 1992; Church et al, 2000; Cotner et al, 1997, 2000; Donachie et al, 2001; Hale et al, 2017; Hoch and Bronk, 2007; Joint et al, 2002; Kirchman, 1990; Kuparinen and Heïnänen, 1993; Liu et al, 2014; Martínez-García et al, 2010a,b; Mills et al, 2008; Obernosterer et al, 2003; Ortega-Retuerta et al, 2012; Pinhassi et al, 2006; Pomeroy et al, 1995; Rivkin and Anderson, 1997; Shiah and Ducklow, 1993; Vadstein, 2011; Yuan et al, 2011; Zweifel et al, 1993). This limitation is widespread across regions of low nutrient concentrations in all major ocean basins (Lennartz et al, 2024). For instance, the degradation rate of fucoidan, a polysaccharide frequently produced by marine brown algae, is strongly dependent on phosphate availability (Xu et al, 2024). In contrast, bacterial uptake is mainly limited by the availability of labile organic carbon in regions where nutrient concentrations (N, P) are high (Church et al, 2000).

Although these nutrient-enrichment experiments yield consistent results, they represent spatially and temporally limited snapshots. High-throughput “omics” technologies are powerful tools to assess basin-scale variations in the microbial metabolic potential (Ustick et al, 2021). Integrating both data sets allows combining mechanistic insights from incubation experiments with the spatial coverage of global genomic surveys. Here, we developed an approach to assess nutrient limitations (phosphorus, nitrogen, iron) for the dominant heterotrophic clade SAR11 based on metagenomes from the Bio-GO-SHIP campaign in a similar approach as previously done for phytoplankton (Ustick et al, 2021). Our metagenomic analysis revealed pronounced macronutrient stress for heterotrophic bacteria in the low- and midlatitude Atlantic and Pacific, except for equatorial upwelling regions (Fig. 1). The geographic pattern of phosphorus and nitrogen limitation are in very good agreement with the most comprehensive study of nutrient enrichment experiments along two Atlantic transects: Therein, phosphorus limits bacterial activity in the North and nitrogen in the South Atlantic, with a transition near 10*^◦^*S (Hale et al, 2017) (Fig. 1). Based on the agreement between metagenomically derived macronutrient stress and nutrient enrichment experiments, we combined the metagenomic indications of both N and P stress to an indicator for macronutrient stress and extrapolated it with a finite-element method (Fig. 1) (Troupin et al, 2012). The underlying rationale for this is that the phosphate concentration in the surface ocean serves as a proxy for macronutrient limitation of microbial carbon uptake (Fig. S3): Macronutrient stress is high largely across low latitudes in the Atlantic Ocean and the subtropical regions of the Pacific (Fig. 1). In the Southern Ocean, in some shelf regions and in the equatorial Pacific, on the other hand, metagenomic data indicate that the heterotrophic microbial community does not face considerable nutrient stress. These observations align with nutrient enrichment experiments that demonstrate carbon limitation in these regions. Taken together, these metagenomic and experimental analyses provide a consistent picture. Multiple lines of evidence including experimental, metagenomic and theoretical approaches suggest macronutrient limitation of heterotrophic microbial activity in oligotrophic regions of the surface ocean.

**Fig. 1:**
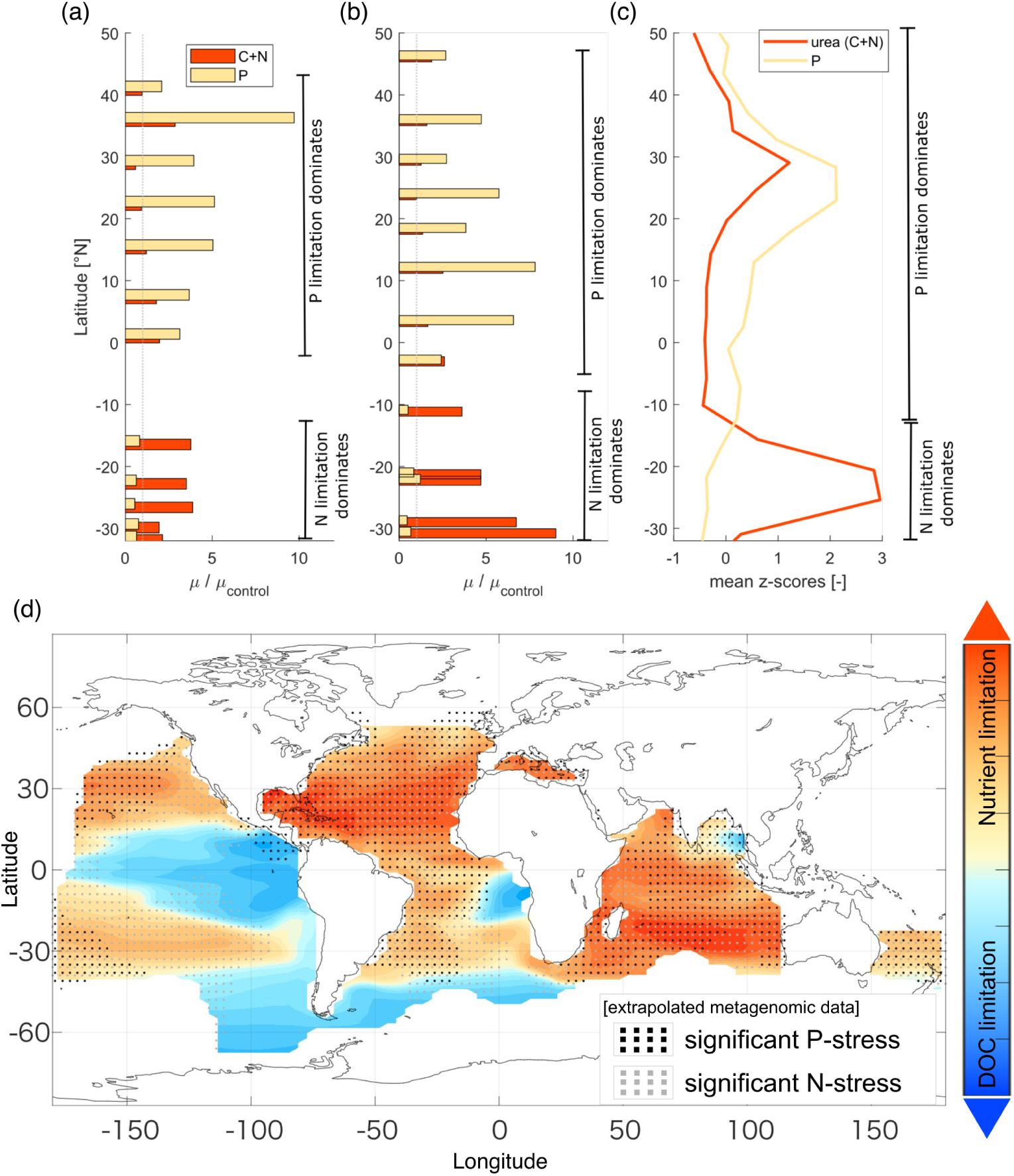
Limiting factors of microbial growth demonstrated by three different methodologies. (a) growth enhancement in an incubation study relative to control upon P and C+N addition for an Atlantic transect (AMT16) in May/June 2005 from Hale et al (2017). The dotted grey line indicates the limitation threshold where growth in nutrient enriched samples equals the growth in the control. (b) same as (a) but for AMT transect 17 in Oct/Nov 2005. (c) nutrient stress derived from metagenomic data from BIO-GOSHIP, plotted as average standard scores (or z-scores) of the relative abundance of the respective nutrient transporters (see methods). In all panels, red indicates nitrogen limitation and yellow indicates phosphorus limitation. Annotations indicate latitudinal range where the respective limitation or stress dominates. (d) Simulated pattern of the limiting factor with the MICDOC-coupled ESM, differentiating between DOC limitation and macronutrient limitation (combining P+N stress in a and b). The growth of heterotrophic bacteria is limited by the availability of DOC in the blue regions, while it is macronutrient-limited (representing N and P limitation) in the orange regions. Interpolated metagenomic data showing macronutrient stresses as hatches on top of the simulated distribution of limiting factors in the model in the background.

### Impacts of nutrient limitation on DOC and DOP

To test the hypothesis of nutrient limitation as a main driver of basin-scale DOC distribution, we modified the existing DOC component (MICDOC, model of microbial DOC interactions) of the University of Victoria Earth System Model of Intermediate Complexity (UVic) (Lennartz et al, 2024). This ESM includes a marine ecosystem module that simulates the interactions among four key state variables, namely nutrient, phytoplankton (diazotrophs and others), zooplankton and detritus (a NPZD model) (Keller et al, 2012; Mengis et al, 2020). Our modifications include two key changes. First, the coupling between the bacteria–DOC component and the biogeochemical host model was made fully bidirectional, allowing for dynamic feedbacks between carbon and nutrient cycles in the model. Secondly, dissolved organic macronutrient (organic phosphorus) was implemented as an additional component, in order to resolve the ability of heterotrophic microorganisms to take up both organic and inorganic nutrients.

In the model, microbial growth–and thus their capacity to degrade DOM–is limited by the availability of DOC or macronutrients (organic and inorganic phosphorus treated as interchangeable P resources) following Liebig’s law of the minimum. In this framework, phosphate concentration serves as a proxy for macronutrient stress. Consistently, the patterns of overall nutrient stress obtained from the model closely resemble those observed for SAR11 in metagenomic data (Fig. 1).

The nutrient limitation of the heterotrophic microbial community directly impacts the turnover and the basin-scale distribution of DOM (i.e. DOC and dissolved organic phosphorus (DOP)). We performed an ensemble of 279 simulations with varying parameter values, and identified the best 20 simulations in terms of reproducing the global distribution of DOC (Hansell et al, 2021) and DOP (Knapp et al, 2022) in the modern surface ocean (Fig. 2; Fig. S5). The growth, hence biomass, of the microbial consumers is determined by the least available resource and thus, the surplus of remaining resources (DOC or DOP) accumulates. This process largely shapes the global distribution of DOM in our model, which is remarkably similar to observations (Fig. 2b; Fig. S5b). Observed zonal-mean DOC concentrations range between ∼45 mmol m*^−^*^3^ and ∼75 mmol m*^−^*^3^. The latitudinal DOC gradient shows differences of 25–30 mmol m*^−^*^3^ between high and low latitudes. This general feature of DOC distribution is well captured by the model simulation. Overall, the water-column DOC profiles simulated by MICDOCv2.0 closely match the comprehensive observational dataset of about 40,000 DOC concentrations across the global ocean (Hansell et al, 2021). The latitudinal gradient as well as the depth gradient are reproduced (Fig. 2). This agreement is even most robust at depths shallower than 1000 m (Fig. 2c; Fig. S4), where macronutrient limitation primarily occurs. Here, globally averaged observations and model simulations agree within <1 mmol m*^−^*^3^. We note that in the abyssal ocean, simulated concentrations are ca. 2 mmol m*^−^*^3^ higher than the average observed concentration, still lying within the overall uncertainty range. These depths, however, are not affected by nutrient limitation, making them independent from our analysis above. The overall root mean square error is 8.2 mmol m*^−^*^3^, which compares well to similar approaches following first-order decay rates of prescribed reactivity fractions (Flanjak et al, 2025; Hansell et al, 2012). In addition, the model is also able to reproduce DOP concentrations (Fig. S5a,b) and the resulting DOC:DOP stoichiometry across the ocean (Fig. 2d; Fig. S5c,d). Especially in regions with high nutrient stress, the DOC:DOP ratio increases from a global average of 200–300 to 300–500, which is reasonably consistent with observation-based data (see supplements).

**Fig. 2:**
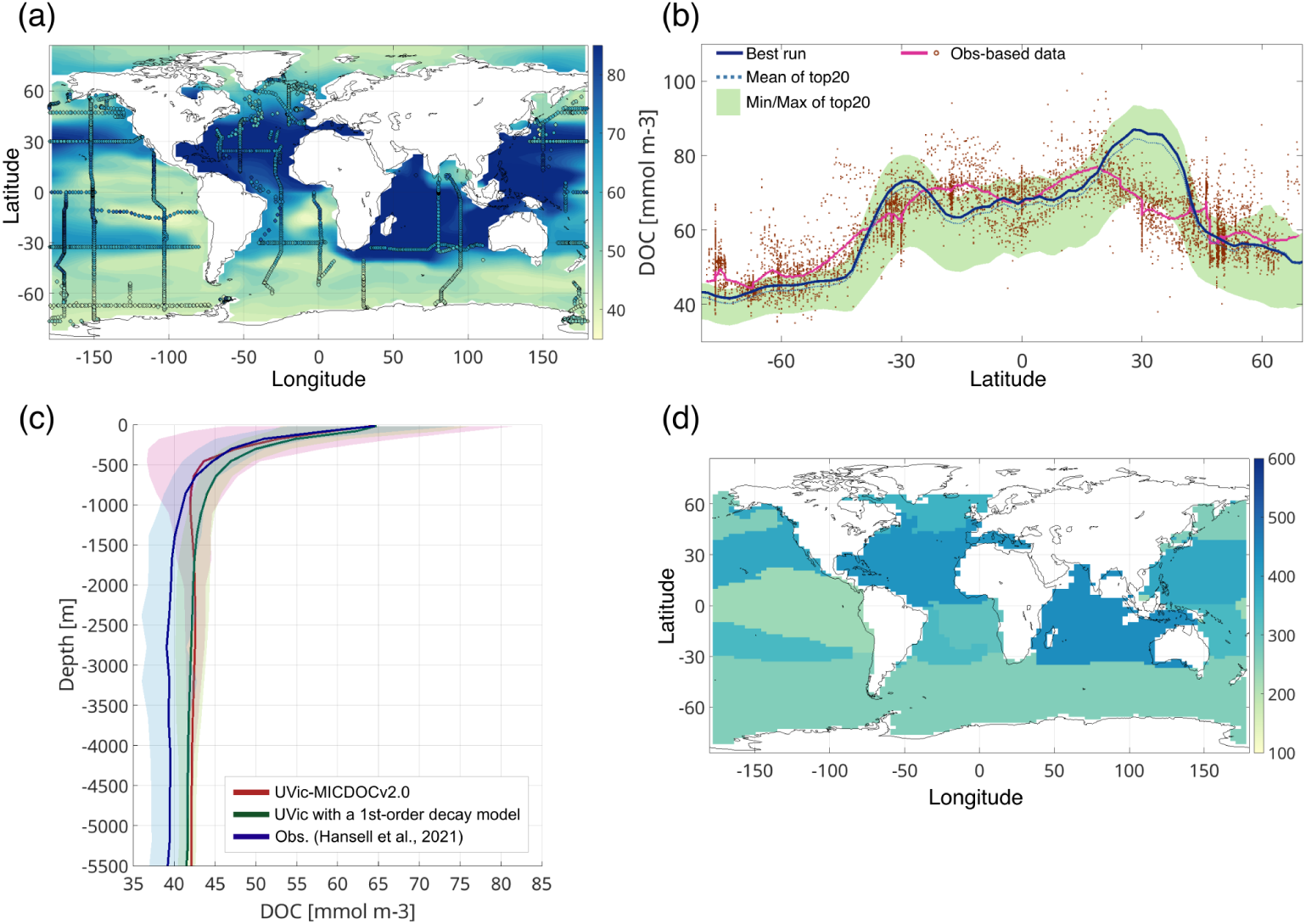
Model performance of the DOC simulations. Concentrations of DOC for the upper 50 m are shown alongside observation-based data as (a) a global map of surface DOC concentrations and (b) as a zonal average (surface). Model results represent the 20-year mean around the year 2000, while observations are from Hansell et al (2021). (b) zonal mean concentrations for the best simulation (blue), the mean of the top 20 runs (blue dotted), and the observations (red). The light-green band denotes the range spanned by the top 20 runs. (c) global mean vertical profiles of DOC concentrations simulated with UVic-MICDOCv2.0 (red), observations (blue) (Hansell et al, 2021), and UVic with a 1st-order-decay model (green). The shaded areas indicate one standard deviation. (d) simulated DOC:DOP stoichiometry in biogeochemically defined regions. As in Liang et al (2023), the region boundaries are defined as 0.3 mmol m*^−^*^3^ of simulated surface phosphate concentration.

### The future fate of the marine DOC inventory

With ongoing warming and the corresponding intensified stratification of the water column, nutrient stress is projected to increase in high-emission future scenarios (e.g. Chavez et al, 2011; Falkowski, 2012). This affects both the DOC production via phytoplankton and the DOC consumption by heterotrophic microorganisms. We carried out transient simulations starting from the pre-industrial (year 1850) state until year 2200 to examine the evolution of the marine DOC reservoir under scenarios of future climate change. The background climate and ocean physics were simulated following the shared socioeconomic pathway (SSP), where we forced the model with a prescribed trajectory of CO_2_ concentration (Meinshausen et al, 2020). In our simulation, DOC export at 100 m depth is 2.1 GtC year*^−^*^1^ under present-day conditions and 2.4 GtC year*^−^*^1^ in year 2200. These values correspond to 19% and 23% of particulate organic carbon (POC) export. The present day rates are consistent with previous estimates (Hansell et al, 2009; Roshan and DeVries, 2017; Wang et al, 2023; Flanjak et al, 2025). This indicates that marine DOC will play a more important role in the biological carbon sink in the future, consistent with the hypothesis of Roshan and DeVries (2017). The impact on carbon sequestration is even more notable in the change of the standing stock of the DOC inventory: The projected size of the DOC reservoir monotonically grows until the end of the year 2200 (Fig. 3a). With the best performing parameter setting, the DOC-based carbon sequestration increases by 39 GtC by 2200 under the highest emission scenario SSP5-8.5. The best 20 simulations comprise a range of an additional storage of 18-44 GtC by 2200.

**Fig. 3:**
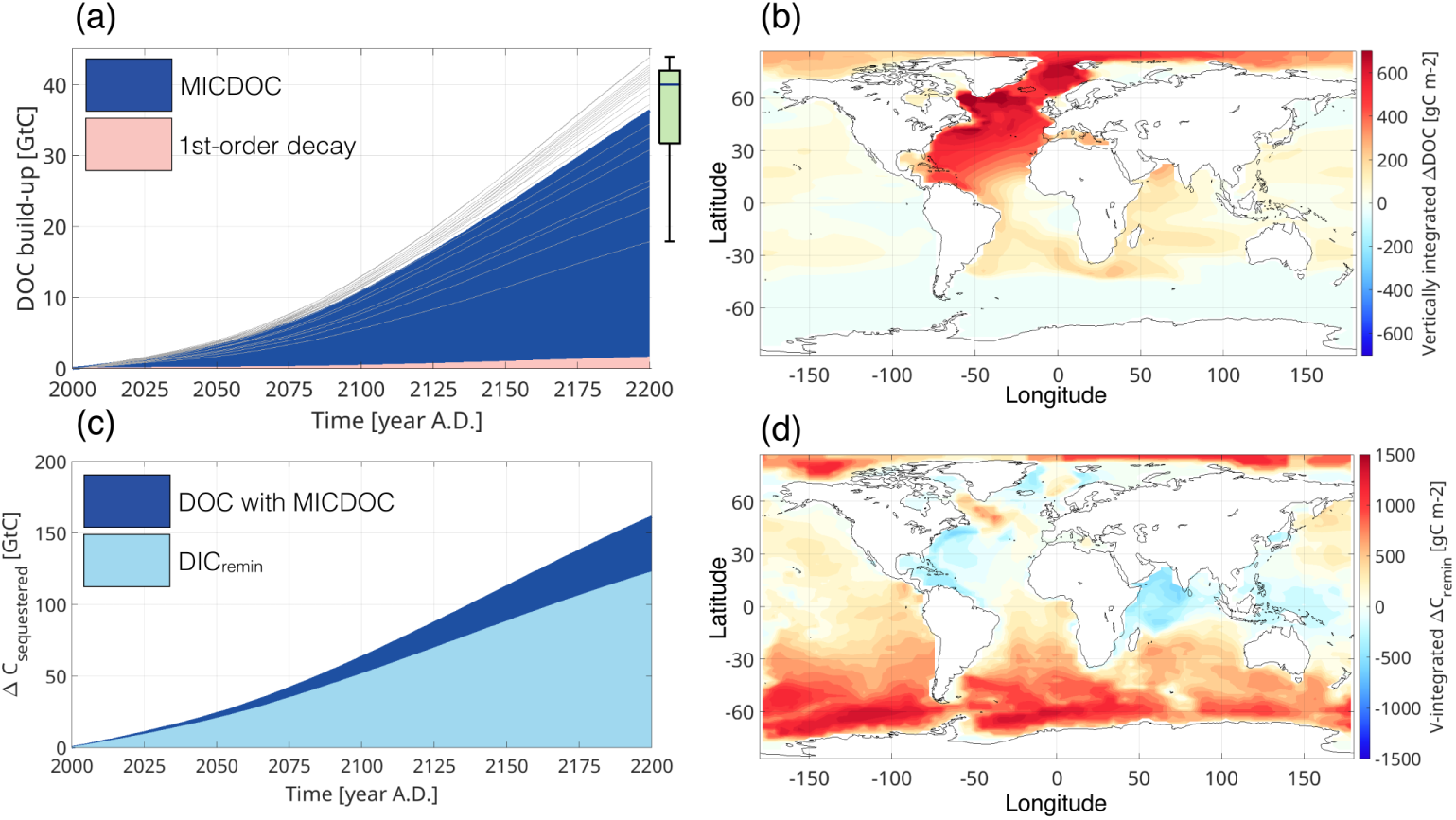
Projected evolution of the DOC pool under a high-emission scenario (SSP5-8.5). (a) Simulated growth of global DOC pool since the year 2000 for the best-performing UVic-MICDOCv2.0 run (blue), for individual top-20 runs (grey lines), and for UVic with a 1st-order decay model based on Hansell et al (2012) (pink). The box plot shows the spread of the top-20 estimates at year 2200. (b) Vertically integrated amount of DOC at year 2200 that has accumulated since year 2000 by the MCP. (c) The increment of global DOC pool as compared with that of carbon sequestered by the BCP (DIC_remin_). (d) Same as in (b), but for DIC_remin_.

This accumulation of DOC augments the biological carbon pump (BCP), the process by which carbon is transported to depth via the sinking of biogenic debris. Here, we define the long-term BCP-based carbon storage as the amount of DIC derived from the remineralization of sinking debris (DIC_remin_) that is stored in the deep ocean 1000 m water depth). DIC_remin_ was estimated from Apparent Oxygen Utilization (AOU) and the stoichiometry of particulate organic matter as done previously in Frenger et al (2024). The amount of DIC_remin_ stored in the deep ocean is also projected to increase in the future due to changes in the ocean circulation (Fig. 3c). The accumulation of DOC as estimated with our model is substantial, increasing the BCP-based carbon sequestration by 30% (Fig. 3c).

Uncertainty remains due to the simplifications and model biases discussed above. However, the amount of the DOC buildup in the “top20” ensemble runs ranges from 18 to 44 GtC. More than half (13 out of 20) of the model runs yield estimates between 37 and 44 GtC, underscoring the robustness of the projected future enhancement of carbon storage mediated by nutrient limitation of bacterial DOC uptake. The positive model bias in the present-day DOC concentrations is minor (<1 mmol m*^−^*^3^) in the 500–2000 m depth range in the Northern Atlantic where a large part of the accumlated DOC is stored until the year 2200 (Fig. 3b).

This feedback has a high potential to impact carbon storage on centennial time scales, with an effect on the same order of magnitude as BCP-driven carbon sequestration. It remains to be tested how other potentially influential factors impact the future inventory size. Future warming and acidification are expected to reshape marine microbial communities (Burrell et al, 2017; Deppeler et al, 2020). These changes include regionally specific shifts in phytoplankton composition, altered microbial diversity and function, and feedbacks on biogeochemical cycles, carbon export and DOC turnover (e.g. Dutkiewicz et al, 2015; Lønborg et al, 2020; Lu et al, 2025). For example, seawater temperature might influence microbial processes involved in DOC degradation (e.g. Lønborg et al, 2018). However, highest DOC concentrations are not found in warmest regions, suggesting nutrient limitation outweighs temperature effects on DOC uptake rates at present.

The mechanism and significance of the future DOC build-up is independent of the emission scenario. We conducted simulations for two other SSP scenarios, namely SSP3-7.0 and SSP2-4.5, with the best-performing parameter set. For those two additional scenarios, the estimated increase in DOC reservoirs by the year 2200 is ∼32 GtC and ∼19 GtC, respectively. On the other hand, the increase in DIC_remin_ below 1000 m is ∼109 GtC and ∼78 GtC. Therefore, the MCP would contribute to the relative enhancement of carbon sequestration to a similar degree (i.e. increase by 25–30%) irrespective of future pathways.

Importantly, the expected change in the surface DOC inventory varies regionally (Fig. 4). In macronutrient-limited regions, reduced macronutrient concentrations decrease primary production and thereby DOC supply. However, this decline is counteracted by reduced DOC remineralization, as bacteria are likewise constrained by macronutrient scarcity. Because the reduction in remineralization exceeds the decrease in supply in our simulations, the net effect leads to a quantitatively significant DOC accumulation (Fig. 4). The relative sensitivity of phytoplankton and heterotrophic bacteria to nutrient availability is supported by microcosm experiments (Martínez-García et al, 2010).

**Fig. 4:**
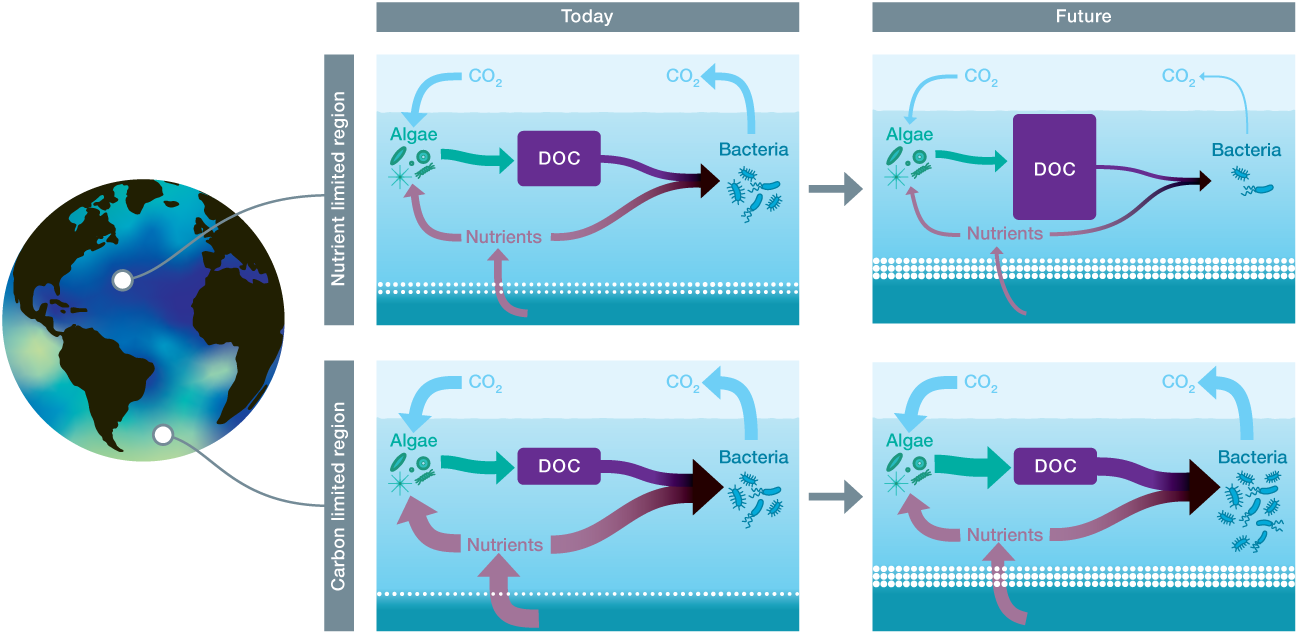
Carbon reservoir size in macronutrient- and carbon-limited oceanic regions: present and future projections. The accumulation of DOC is defined by the balance between DOC production by algae and DOC consumption by bacteria. Since macronutrients and DOC are taken up in specific ratios, the nonlimiting component accumulates. In oligotrophic regions today, where nutrient availability is low, DOC accumulates, whereas lower DOC concentrations are found when DOC is limiting. In a high-emission future scenario, increasing stratification is expected to reduce nutrient supply to surface waters, impacting both algal DOC production and bacterial consumption. Despite declining algal production, oligotrophic regions are projected to experience an increase in the DOC reservoir due to reduced bacterial uptake under nutrient limitations. In contrast, in carbon-limited regions, the overall DOC inventory is stable against variation in nutrient concentrations, because bacterial uptake is limited by the DOC concentration.

In the DOC-limited regions, DOC supply increases are offset by a similar amount of increases in bacterial consumption (Mentges et al, 2019, 2020; Dittmar et al, 2021) and DOC concentrations only show minimal response to environmental change (Fig. 4). Given global nutrient distributions and the projected expansion of oligotrophic regions, mean surface DOC is expected to rise overall in the future. Globally, the vertically integrated amount of the increase in DOC between year 2000 and 2200 is largest in the North Atlantic. This pattern arises from the transport of DOC-rich surface waters by deep convection in the high-latitude North Atlantic, followed by their subsequent spread across the deep North Atlantic (Fig. 3b).

### Mechanisms behind the DOC accumulation

For the present day, the water-column DOC profile above 1000 m simulated with MICDOCv2.0 closely matches that of extensive observations (Hansell et al, 2021) (Fig. 2c; Fig. S4). An additional simulation, employing a first-order decay model instead of explicit microbial consumers, exhibits a similar profile across most depths. However, the future projection with MICDOCv2.0 differs from those derived from the traditional first-order decay model that does not explicitly represent microbial DOC degradation. Importantly, because MICDOCv2.0 resolves the underlying processes rather than relying on apparent net removal rates, it enables separate evaluation of environmental controls on both DOC production and degradation–an essential requirement for projecting future changes in both processes.

The projected DOC accumulation until the year 2200 differs by 16–42 GtC between the two model versions, with MICDOCv2.0 yielding substantially greater carbon storage than the model that extrapolates net removal rates. In the UVicESCM simulation with first-order decay, DOC accumulation was only ∼1.6 GtC (Fig. 3a). Another net-removal-rate-based model projected a ∼6 GtC global DOC increase between 2000–2200 driven largely by higher primary production (Gilchrist, 2022).

This difference in storage potential arises from the dynamic feedback between carbon and nutrient cycles that is only represented in the MICDOC model. In the conventional net-removal-rate approach, first-order decay rates for DOC remineralization are fixed and do not respond to changing environmental conditions. Moreover, the prescribed lifetimes of the three DOC fractions (1.5, 20, and 16000 years, following Hansell et al (2012)) represent net removal rates chosen to maintain a DOC concentration of approximately 35 mmol m*^−^*^3^ in the oldest water masses, effectively pre-imposing an inert DOC behavior on the model.

In contrast, DOC uptake by heterotrophic microbes in MICDOC responds dynamically to DOC and macronutrient concentrations. A key driver of the simulated DOC increase is the decline in surface macronutrient levels by the year 2200 relative to today (Fig. S6), caused partly by reduced nutrient supply from depth under warming-induced stratification and partly by intensified macronutrient consumption linked to temperature-driven increase in maximum primary production rates that is implemented in UVicESCM (Keller et al, 2012).

Our simulations demonstrate that the DOC–macronutrient feedback driving a substantial increase in DOC storage is not captured by traditional modelling approaches. This highlights the need to assess environmental controls on production and consumption separately, rather than combining them into a single net removal rate that does not respond to changing environmental conditions. A process-oriented, mechanistic representation of remineralization is therefore essential to understand the feedbacks in the marine carbon cycle.

### Implications for DOC in a changing climate

This study presents the first future projections of the bulk marine DOC pool using a fully coupled DOM–ocean model that explicitly resolves bacterial remineralization of DOC and accurately captures present-day DOC distributions. The model reproduces key global features, including present-day DOC concentrations, stoichiometry, and the spatial patterns of microbial nutrient limitation. Under a high-emission climate scenario, it predicts substantial DOC accumulation–particularly in the North Atlantic–driven by intensified stratification and nutrient stress in heterotrophic microbial communities.

These findings challenge the prevailing view of the DOC reservoir as a static and inert component of the carbon cycle. Instead, they reveal a system responsive to nutrient dynamics and climate changes, with the DOC pool increasing by up to 2 PgC per decade. Projected accumulations of 18–44 PgC by the year 2200 exceed the blue carbon sequestration potential of vegetated coastal ecosystems by an order of magnitude (Bertram et al, 2021), and represents a ∼30% enhancement of the biological carbon pump. Our results underscore the need to incorporate microbial physiology and nutrient-driven feedbacks into Earth system models to improve projections of ocean carbon storage. The consistent emergence of nutrient-stress-driven DOC accumulation across our ensemble simulations provides robust evidence for a negative feedback on marine carbon storage under a future climate scenario. In addition to their response to nutrient supply, concurrent warming and acidification induce complex, often non-linear changes in microbial community structure and function (e.g. O’Brien et al, 2016). The responsiveness of the DOC pool on decadal to centennial timescales underscores the need to better understande future microbe–DOC interactions under changing environmental conditions.

Integrating multi-omics, experimental, and modelling approaches offers a path toward more mechanistic assessments of marine carbon cycling. With ongoing and emerging ocean observing initiatives such as BioGeoScapes (e.g. Tagliabue, 2023), Bio-GO-SHIP (e.g. Clayton et al, 2022), and Tara (e.g. Pesant et al, 2015), enhanced model–data integration will further refine projections of ocean carbon sequestration and support science-based marine carbon management. Here, we provide a platform that enables a process-oriented implementation of a holistic view of the carbon cycle, which explicitly includes the environmental controls on the remineralization of DOC back to CO_2_. The contrasting outcomes of our two model versions–one based on net removal rates and one grounded in ecological mechanisms–demonstrate the importance of explicitly representing the DOC pool in future climate scenarios and carbon storage strategies.

## Acknowledgements

We acknowledge funding by the Ministry for Science and Culture of the State Lower-Saxony (MWK), granted to STL (Grant 16TTP079)).

## Author contribution

TKN and STL developed MICDOC version 2. TKN and STL designed the experiments, and TKN carried them out. ACM provided the metagenomic data and analysis. STL and TD conceptualized the overarching research project. All authors interpreted and discussed the model outputs. TKN prepared the manuscript with contributions from all co-authors.

