## Supplementary information for "Enhanced carbon storage in dissolved organic matter in a future oligotrophic ocean"

### Methods

#### The microbial-DOC model MICDOCv2.0

We employed an Earth system model of intermediate complexity (EMIC) for the global carbon cycle that is coupled with a dynamical DOM–microbes–nutrients system (MICDOCv2.0, Fig. S1). MICDOC depicts the resource–consumer interactions among DOC, DOP, microbes, and macronutrients (Tilman, 1982). The newly developed coupled model allows channeling of carbon fixed by primary production into different routes of export, i.e. POC or DOC, to assess its feedback on the global carbon and nutrient cycles.

For the EMIC, the University of Victoria Earth System Climate Model version 2.10 (UVic ESCM 2.10; Mengis et al (2020)) was employed, which provides 3-dimensional ocean physics and NPZD-based marine biogeochemistry (Keller et al, 2012). The ocean component had the horizontal resolution of 3.6° in longitude and 1.8° in latitude, and 19 depth levels.

To assess the DOM–microbes–nutrients dynamics, we substantially extended the existing version of our model (MICDOC version 1; Lennartz et al (2024)). The main differences from the previous version is twofold: First, the UVic ESCM and the MICDOC model is now interactively coupled, so that DIC concentration and macronutrient concentrations of the UVic ESCM host model are affected by MICDOC dynamics and vice versa. Second, the DOM in the MICDOC model has two separate components of DOC and DOP, allowing us to assess the impact of organic nutrients for heterotrophic microbes. The microbial component follows a prescribed P:C ratio ($\theta$), while the host UVic model has a separate stoichiometric ratio ($\varphi$) for the other biogeochemical components.

The temporal evolution of the concentrations of heterotrophic microbes, DOC, and DOP is calculated using the following ordinary differential equations.

$$\frac{d[\mathrm{MIC}]}{dt} = \eta \cdot docu - docl \tag{1}$$

$$\frac{d[\mathrm{DOC}]}{dt} = docs + docl + docr - docu \tag{2}$$

$$\frac{d[\mathrm{DOP}]}{dt} = \varphi \cdot docs + \theta \cdot docl + \frac{\mu_{\mathrm{DOP}}}{\mu_{\mathrm{DOP}} + \mu_{\mathrm{PO4}}} \cdot \theta \cdot docr - \frac{\mu_{\mathrm{DOP}}}{\mu_{\mathrm{DOP}} + \mu_{\mathrm{PO4}}} \cdot \theta \cdot docu \tag{3}$$

The uptake of DOC by heterotrophic microbes (*docu*) is limited by the availability of DOM and macronutrients (both organic and inorganic) with the following uptake kinetics:

$$docu = \rho \cdot \min(\mu_{\mathrm{DOC}}, (\mu_{\mathrm{DOP}} + \mu_{\mathrm{PO4}})) \cdot [\mathrm{MIC}] \tag{4}$$

where $\rho$ is the potential maximal growth rate of microbes, [MIC] is the local concentration of heterotrophic microbes, and $\min(\mu_{\mathrm{DOC}}, (\mu_{\mathrm{DOP}} + \mu_{\mathrm{PO4}}))$ is a limiting factor that determines the actual growth rate under a given environment. The limiting factor of growth is determined in a "law of the minimum" relationship, in which the uptake rates of DOC and phosphorus (i.e. the sum of DOP and DIP) are compared and the overall uptake is determined by the lowest of the two fluxes. The uptake rate of the nonlimiting resource is scaled with the P:C ratio of microbial biomass ($\theta$). In this model, we set phosphorus as representative of macronutrients, because both N and P concentrations show globally similar distributions. Out of the DOM uptake, a fraction $\eta$ is fixed into microbial biomass.

The potential uptake rates of each substrate/nutrient ($\mu_{\mathrm{i}}$, with i being DOC, DOP or $PO_4$) is a function of its concentration in the seawater following Monod kinetics as follows:

$$\begin{cases} \mu_{\mathrm{DOC}} = n_{uc} \dfrac{[\mathrm{DOC}]}{\kappa_{\mathrm{DOC}}} \\ \mu_{\mathrm{DOP}} = n_{up} \dfrac{[\mathrm{DOP}]}{\kappa_{\mathrm{DOP}}} \\ \mu_{\mathrm{PO4}} = \dfrac{[\mathrm{PO_4}]}{[\mathrm{PO_4}] + \kappa_{\mathrm{PO4}}} \end{cases} \tag{5}$$

where $n_{uc}$ and $n_{up}$ are the ratio of dissolved organic matter that is taken up by an average heterotrophic microbe, and $\kappa_{\mathrm{i}}$ are the half-saturation constants. For DOC and DOP, the approach by Mentges et al (2020) was adopted, which effectively lowers the maximum growth rate for organic matter to account for the diversity of organic compounds, from which only a subset is available to a single bacterium. This approach has been derived in Mentges et al (2020) and tested in Owusu et al (2026) against the original, detailed network box model that MICDOC is build on (Mentges et al, 2019).

The lysis of heterotrophic microbes (*docl*) is proportional to the mortality rate $\lambda$ and is assumed to contribute to a source of DOC.

$$docl = \lambda \cdot [\mathrm{MIC}] \tag{6}$$

The fraction of DOM that is released back to the DOM pools (*docr*) is given by

$$docr = \beta(1 - \eta) \cdot docu \tag{7}$$

The remaining fraction of DOM, $(1 - \beta)(1 - \eta)$, is respired and transferred to the dissolved inorganic carbon pool in the host biogeochemical model. The newly developed model allows channeling of carbon and nutrients fixed by primary production into different routes of export, i.e. POM or DOM. The proportion that goes to the dissolved component is given by a parameter $\gamma$, which determines the supply of DOC (*docs*) together with the production of detritus modeled by the UVic biogeochemistry.

The coupled UVic–MICDOC model was tuned so that the model–data mismatch defined as the root mean square error (RMSE) was minimized. The tuning target was given by the observation-based dataset for the distribution of DOC (Hansell et al, 2021) and DOP (Knapp et al, 2022) in the global ocean. For the model–data comparison, data points in open ocean were only taken into account, considering that characteristic processes in coastal and arctic regions (e.g. riverine inflow of biogeochemical matter or small-scale shelf processes) were not explicitly modeled in our framework.

The coupled model was initialized with a spun-up UVic ESCM state (Mengis et al, 2020) for the conventional UVic variables, and with a homogeneous distribution of MICDOC tracers. We carried out a set of ten-thousand-year model runs with a constant external forcing corresponding to the pre-industrial condition, so that we obtained a quasi-steady model state for each. The ensemble included 250 members, with parameter sets following a Latin hypercube sampling (LHS) design to fill the parameter space with a comparatively smaller number of experiments (Williamson, 2015). We performed additional 29 runs for further manual tuning, summing up to 279 model runs in total. From those, we picked the top 20 individual runs in terms of the model performance (RMSE), for each of which we conducted an subsequent transient run to simulate the time evolution of the system from AD1850 to 2200 with prescribed $CO_2$-concentrations. The parameter set for the best-performing experiment is provided in Table S1. For the climatic forcing of the transient runs, we used $CO_2$ concentrations corresponding to the highest emission scenario (SSP5-8.5) in the shared socio-economic pathway (SSP; Meinshausen et al (2020)).

Table S1 Parameter set of MICDOCv2.0 for the best-performing run

| Parameters | Values | Note |
|---|---|---|
| $\theta$ | 0.01584 | P:C in the microbial biomass [–] |
| $\eta$ | 0.2411 | fraction of uptake converted to micorbial biomass [–] |
| $\rho$ | 1.0 | maximum uptake rate by microbes [$day^{-1}$] |
| $n_{uc}$ | 0.032186 | uptake ratio of DOC [–] |
| $\kappa_{DOC}$ | 7.73 | half saturation constant for DOC [mmol $m^{-3}$] |
| $n_{up}$ | 0.118076 | uptake ratio of DOP [–] |
| $\kappa_{DOP}$ | 0.1444 | half saturation constant for DOP [mmol $m^{-3}$] |
| $\kappa_{PO4}$ | 13.19 | half saturation constant for $PO_4$ [mmol $m^{-3}$] |
| $\lambda$ | 0.042866 | microbial mortality rate [$day^{-1}$] |
| $\beta$ | 0.12 | fraction of uptake released [–] |
| $\gamma$ | 0.15 | production ratio of DOM [–] |

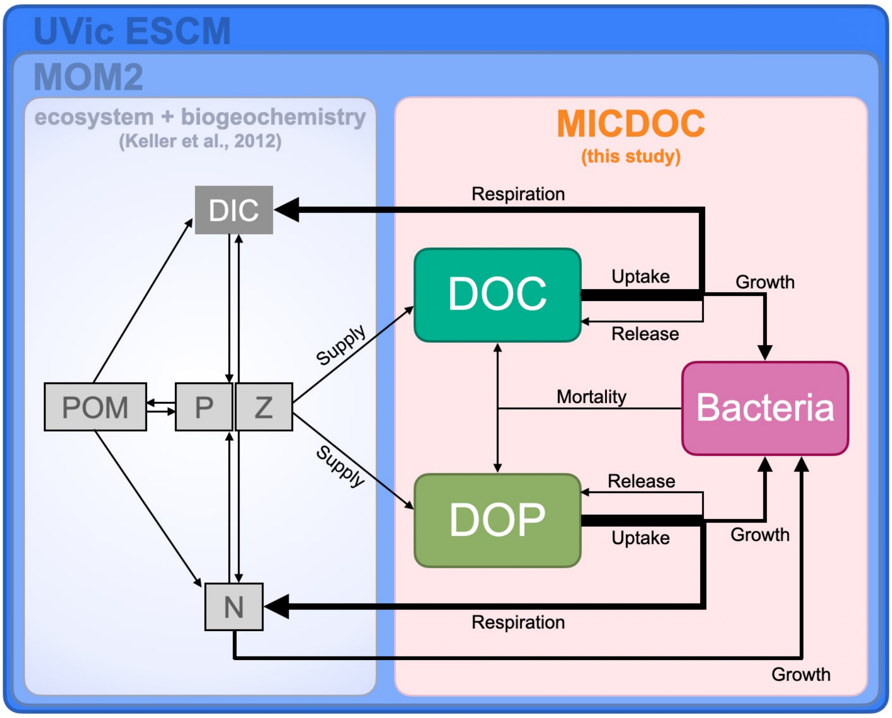


Fig. S1 Schematic of the coupled UVic–MICDOCv2.0 model used in this study. The nutrient–phytoplankton–zooplankton–detritus (NPZD) ecosystem module embedded in UVic ESCM (Keller et al, 2012) is now interactively coupled with the MICDOC model we developed. A fraction of the fixed carbon is channelled into either particulate organic matter (POM) or DOM according to a prescribed ratio, with part of the POM subsequently remineralized into DOM or DIC. The structure of MICDOC is similar to that of the previous version (Lennartz et al, 2024), except for the explicit DOP pool and the interactive flows of C and P between the two modules.

#### DOC modelled with first-order decay

To compare the future simulation with the interactive MICDOCv2.0 model, we contrast this model with a traditional net-removal rate model. This net removal rate type is commonly used in major climate models (Anderson et al, 2015). Here, we use the formulation of Hansell et al (2012). DOC is produced as a fraction of net primary production (NPP), and separated into three different reactivity fractions: semi-labile, semi-refractory and refractory. The reactivity is determined by the first-order rate constant. All three DOC pools are modelled with the following equation:

$$\frac{d\mathrm{DOC}_i}{dt} = s_i \cdot \sqrt{\mathrm{NPP}} - k_i \cdot [\mathrm{DOC}_i] \tag{8}$$

with $s$ being the fraction of NPP channelled into the respective pools $i$ (i.e. semi-labile, semi-refractory, and refractory), and $k$ being the rate constant of the first-order decay [$s^{-1}$]. The parameters for $s$ and $k$ are listed in Table S2. Considering the long lifetime of the refractory fraction, the model was spun up for 124,800 years.

Table S2 Parameters used in the first-order decay model

| Parameters | Values | Note |
|---|---|---|
| $s_1$ | $1.2 \times 10^{-4}$ | semi-labile [(mmol $m^{-3}$ $s^{-1}$)$^{1/2}$] |
| $s_2$ | $3.6 \times 10^{-5}$ | semi-refractory [(mmol $m^{-3}$ $s^{-1}$)$^{1/2}$] |
| $s_3$ | $1.9 \times 10^{-6}$ | refractory [(mmol $m^{-3}$ $s^{-1}$)$^{1/2}$] |
| $k_1$ | $2.114 \times 10^{-8}$ | semi-labile [$s^{-1}$] |
| $k_2$ | $1.586 \times 10^{-9}$ | semi-refractory [$s^{-1}$] |
| $k_3$ | $1.982 \times 10^{-12}$ | refractory [$s^{-1}$] |

#### Nutrient stress derived from metagenomes

Metagenomic data from Bio-Go-Ship campaigns was used to determine nutrient stress, defined here as an enrichment in transporters for N and P relative to the mean content of transporters across all samples. We set Pelagibacter as a representative of nutrient stress in heterotrophic microorganisms, which is a common and ubiquitous heterotrophic microorganism in the ocean (Morris et al, 2002). This approach is conservative, similar to using Prochlorococcus as a representative of phototrophic microorganisms in Ustick et al (2021), because these organisms are among the smallest and, hence, likely least sensitive to nutrient stress. Following Ustick et al (2021), we mapped metagenomic sequences to 37 Pelagibacter genomes representing the major clades (see Table S3 in Ustick et al (2021)). We base this analysis on 21 genes associated with P stress (Table S3), covering inorganic uptake, stress regulation, and use of dissolved organic phosphate (DOP) (Rusch et al, 2007; Coleman and Chisholm, 2010; Martiny et al, 2011). We used 6 genes associated with urea uptake to represent N stress.

As described in Ustick et al (2021), raw reads were quality controlled and adapter sequences were removed using Trimmomatic v0.35. Sequences were mapped using Bowtie2 v2.2.7 against reference genomes with representatives of Pelagibacter as well as other major linages (Prochlorococcus, Synechococcus, and Roseobacter) to help reduce false recruitments. For bowtie 2, we used these flags: –no-unal –local -D 15 -R 2 -L 15 -N 1 –gbar 1 –mp 3. Output files were sorted and indexed using 'samtools' v1.3 into BAM files. Anvi'o v5 was used to profile the recruited reads. All open reading frames were aligned and clustered using NCBI BLAST and MCL through the Anvi'o pangenomic workflow. These clusters were curated and selected by searching for target nutrient stress genes in Matlab. Nutrient stress is calculated as the z-scores of the gene abundances, indicating enrichment of nutrient-stress associated genes above the mean for higher (positive) values.

In order to compare metagenomic stress with model results effectively, the discrete scores of the aggregated P- and N-stress data for each sample location (Fig. S2) were interpolated with the Data-Interpolating Variational Analysis (DIVA) (Troupin et al, 2012) with a correlation length of 20 degrees. Regions with significant nutrient stress defined by a threshold standard score of $-0.5$ gave an excellent agreement with modelled geographic patterns of nutrient stress (Fig. 1d).

Table S3 Clusters of Orthologous Genes (COG IDs) of metagenomic data for nutrient stress used in this study.

| | COG ID | Name |
|---|---|---|
| Phosphorus | COG0226 | ABC-type phosphate transport system, periplasmic component |
| | COG0248 | Exopolyphosphatase/pppGpp-phosphohydrolase |
| | COG0394 | Protein-tyrosine-phosphatase |
| | COG0573 | ABC-type phosphate transport system |
| | COG0581 | ABC-type phosphate transport system |
| | COG0704 | Phosphate uptake regulator |
| | COG0855 | Polyphosphate kinase |
| | COG1117 | ABC-type phosphate transport system, ATPase component |
| | COG2062 | Phosphohistidine phosphatase SixA |
| | COG2908 | UDP-2,3-diacylglucosamine pyrophosphatase LpxH |
| | COG3221 | ABC-type phosphate/phosphonate transport system periplasmic component |
| | COG3454 | Alpha-D-ribose 1-methylphosphonate 5-triphosphate diphosphatase PhnM |
| | COG3624 | Alpha-D-ribose 1-methylphosphonate 5-triphosphate synthase subunit PhnG |
| | COG3625 | Alpha-D-ribose 1-methylphosphonate 5-triphosphate synthase subunit PhnH |
| | COG3626 | Alpha-D-ribose 1-methylphosphonate 5-triphosphate synthase subunit PhnI |
| | COG3627 | Alpha-D-ribose 1-methylphosphonate 5-phosphate C-P lyase |
| | COG3638 | ABC-type phosphate/phosphonate transport system, ATPase component |
| | COG3639 | ABC-type phosphate/phosphonate transport system, permease component |
| | COG4107 | ABC-type phosphonate transport system |
| | COG4778 | Alpha-D-ribose 1-methylphosphonate 5-triphosphate synthase subunit PhnL |
| | COG5002 | Signal transduction histidine kinase |
| Nitrogen | COG0804 | Urease alpha subunit |
| | COG0829 | Urease accessory protein UreH |
| | COG0830 | Urease accessory protein UreF |
| | COG0831 | Urease gamma subunit |
| | COG0832 | Urease beta subunit |
| | COG2371 | Urease accessory protein UreE |

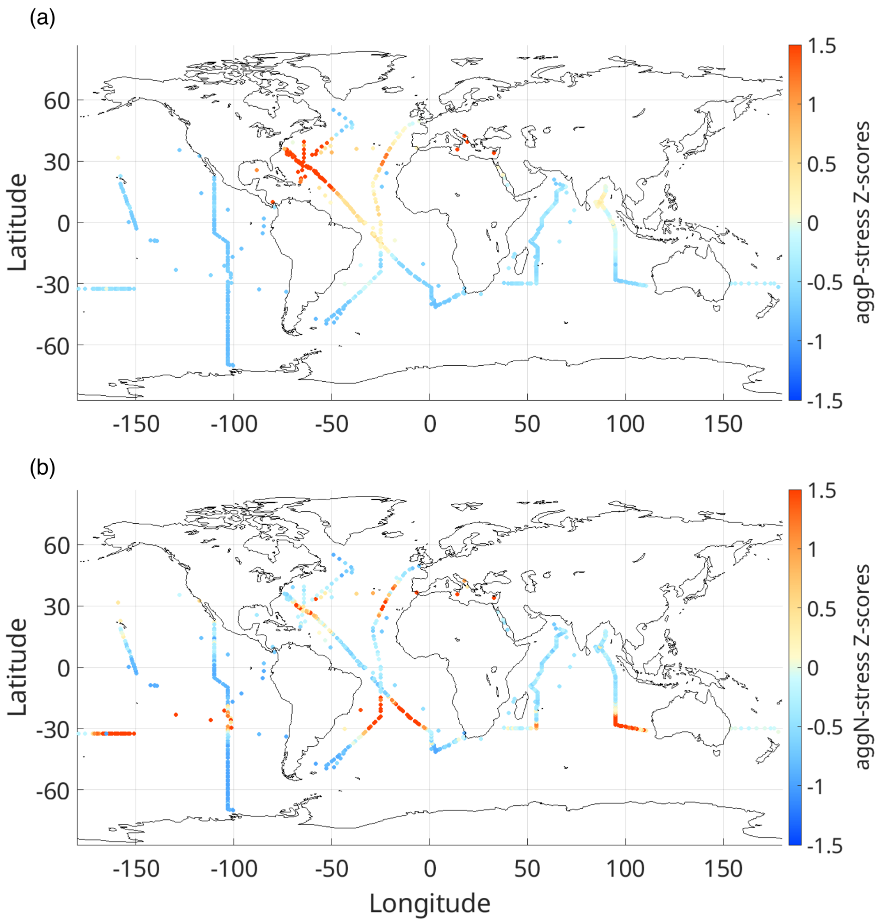


Fig. S2 Sampling locations of the metagenomic datasets used in this study (Bio-Go-Ship, (Larkin et al, 2021)). Normalized gene abundances indicating the severity of nutrient stress for (a) phosphorus and (b) nitrogen are displayed using a color scale.

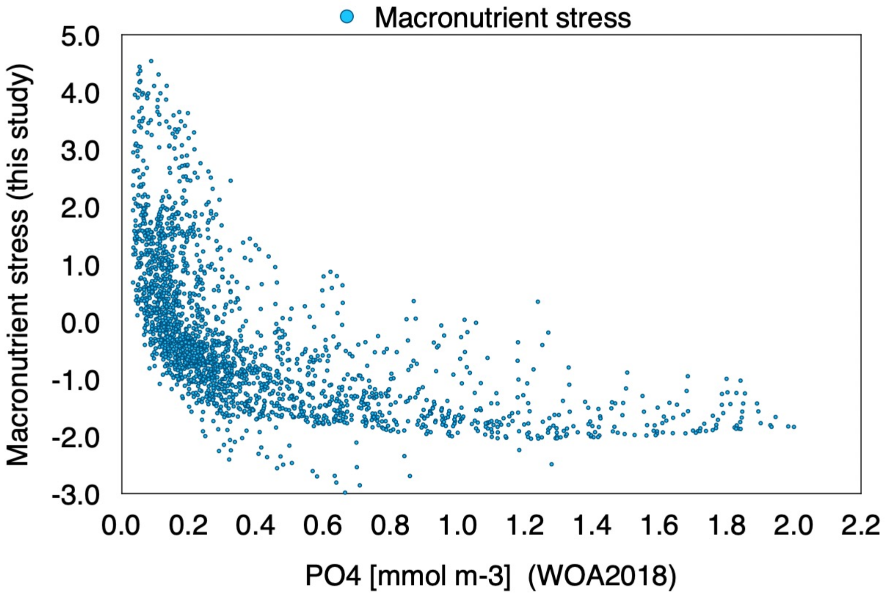


Fig. S3 Correlation between observed phosphate concentrations in the surface ocean (Garcia et al, 2019) and macronutrient stress estimated by the metagenomic data (this study). The macronutrient stress is defined as combined metagenomic indications of both N and P stress by means of their respective z-scores.

### Supplementary text

#### Model performance

For the dissolved organic phosphorus (DOP) concentration, the overall range of concentrations is well reproduced, with a observed and modelled mean concentration of DOP around ∼0.2 mmol $m^{-3}$ (Fig. S5b). Both, model and observations, lack a clear latitudinal structure. The stoichiometry of the modelled DOM, i.e. the regional distribution of the DOC:DOP ratio in the surface ocean, is also consistent with the observation-based estimation within the data uncertainties (Liang et al, 2023) (Fig. S5c,d).

In this study, our focus is on large-scale (basin-scale or greater) features, where the spatial structure of dissolved organic carbon and phosphorus is shaped by the combined influences of biological and physical processes. MICDOCv2.0 demonstrates skill to reproduce such large-scale patterns of DOC distribution, i.e. the latitudinal gradient and accumulation in subtropical gyres. Globally, the overall profile of DOC concentrations is in strong agreement with the observational dataset (Fig. 2c). Small systematic deviations remain, i.e. i.e. positive biases in the nutrient concentrations in upwelling regions are associated with negative biases in the DOC concentration, as discussed in Lennartz et al (2024). The simulation with the lowest RMSE (blue line in Fig. 2b) shows a overestimation of DOC values in the Northern hemisphere around 30°N. However, our assessment is based on the 20 simulations embracing the majority of observed DOC concentrations, ensuring the robustness of our findings.

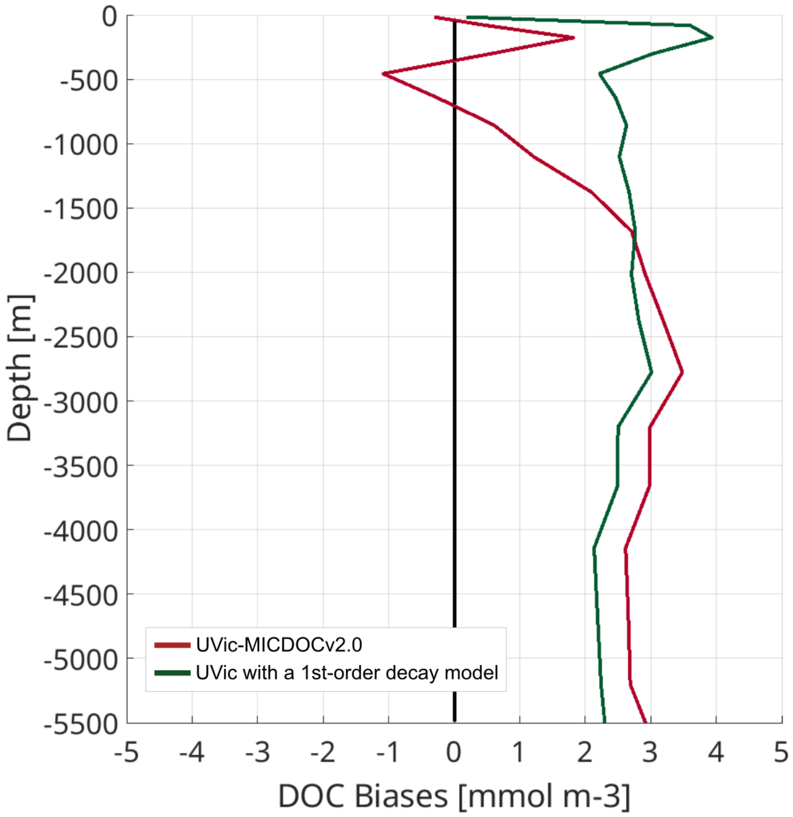


Fig. S4 Deviations in global mean vertical profiles of DOC between the model-based reconstructions (red: UVic-MICDOCv2.0; green: UVic with a 1st-order decay model) and the observations (Hansell et al, 2021).

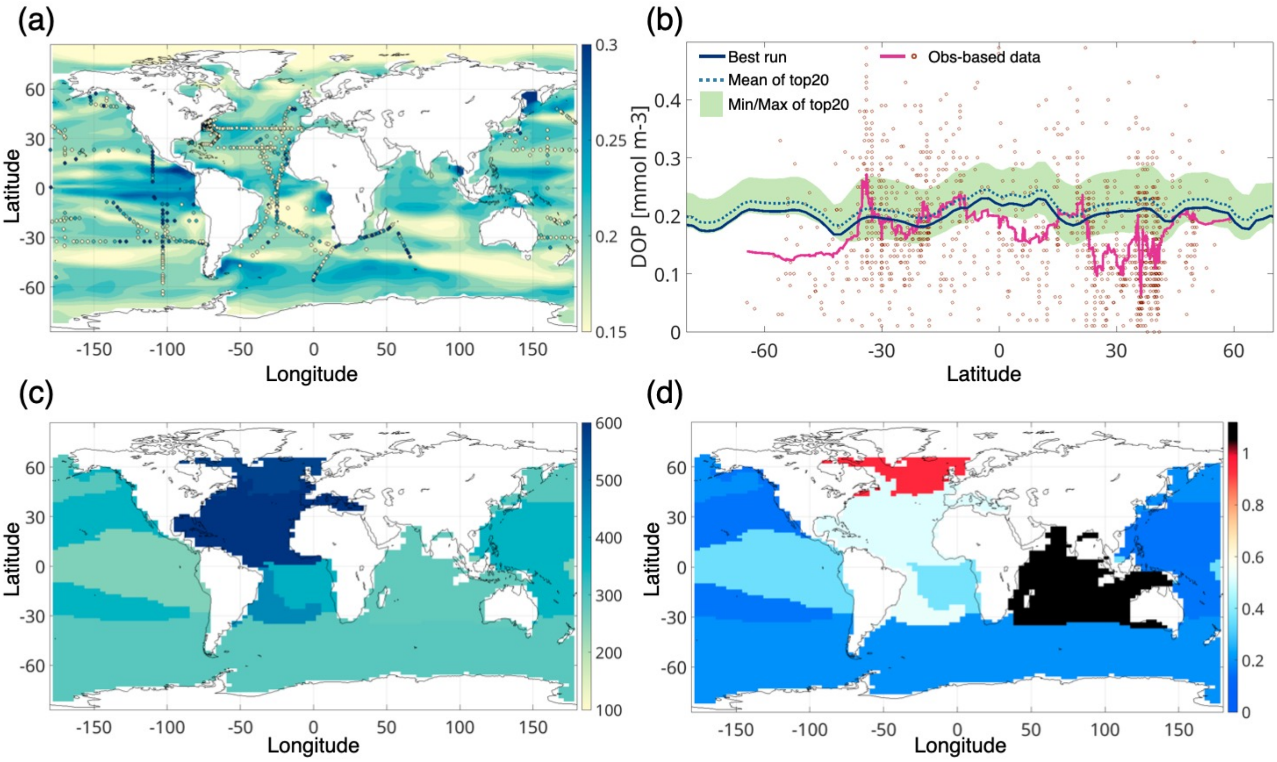

Fig. S5 Simulated distributions of DOP. The concentration for the top 50 m is shown with corresponding observational data in (a) as a global map and (b) as a zonal mean plot. For the model results, the 20-yr average at the year 2000 is presented, with the observations from Knapp et al (2022). In (b), the zonal mean concentrations are shown for the best run (blue), the mean of the top 20 runs (blue dotted), and the observation (red). The light-green band indicates the range covered by the top 20 runs. (c) Observation-based DOC:DOP by Liang et al (2023) allotted to the biogeochemically defined regions according to the modelled nutrient distribution. (d) Model–data mismatch in DOC:DOP that is normalized by the data uncertainties (i.e. abs(model — data)/uncertainty). In the regions where the value is less than 1, the simulated stoichiometry is consistent with the observation within the respective uncertainty.

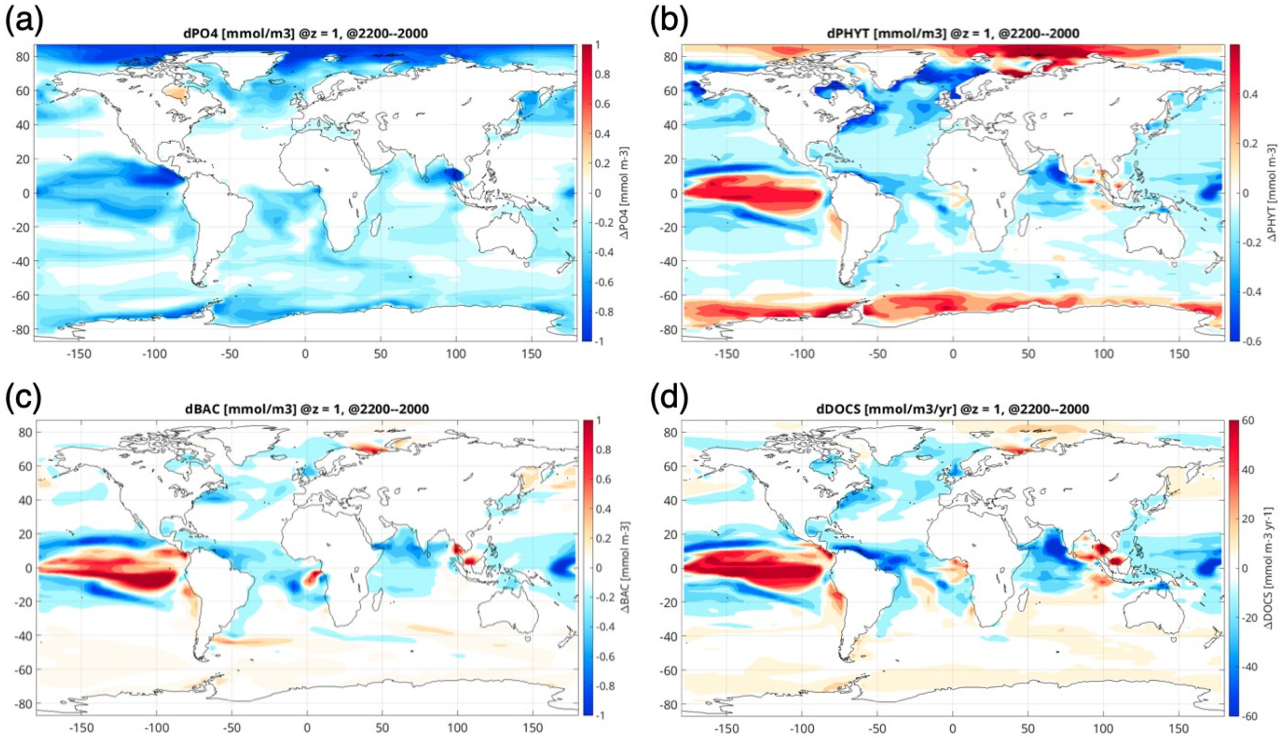


Fig. S6 Projected changes in the surface concentration of various biogeochemical tracers in the transient future simulation with the best-performing parameter set: (a) phosphate, (b) phytoplankton biomass, (c) bacterial biomass, and (d) DOC supply. The differences between AD2200 and AD2000 are shown.
